## Supplemental Figures for "*map3k1* is required for spatial restriction of progenitor differentiation in planarians"

### Figure 1- figure supplement 1

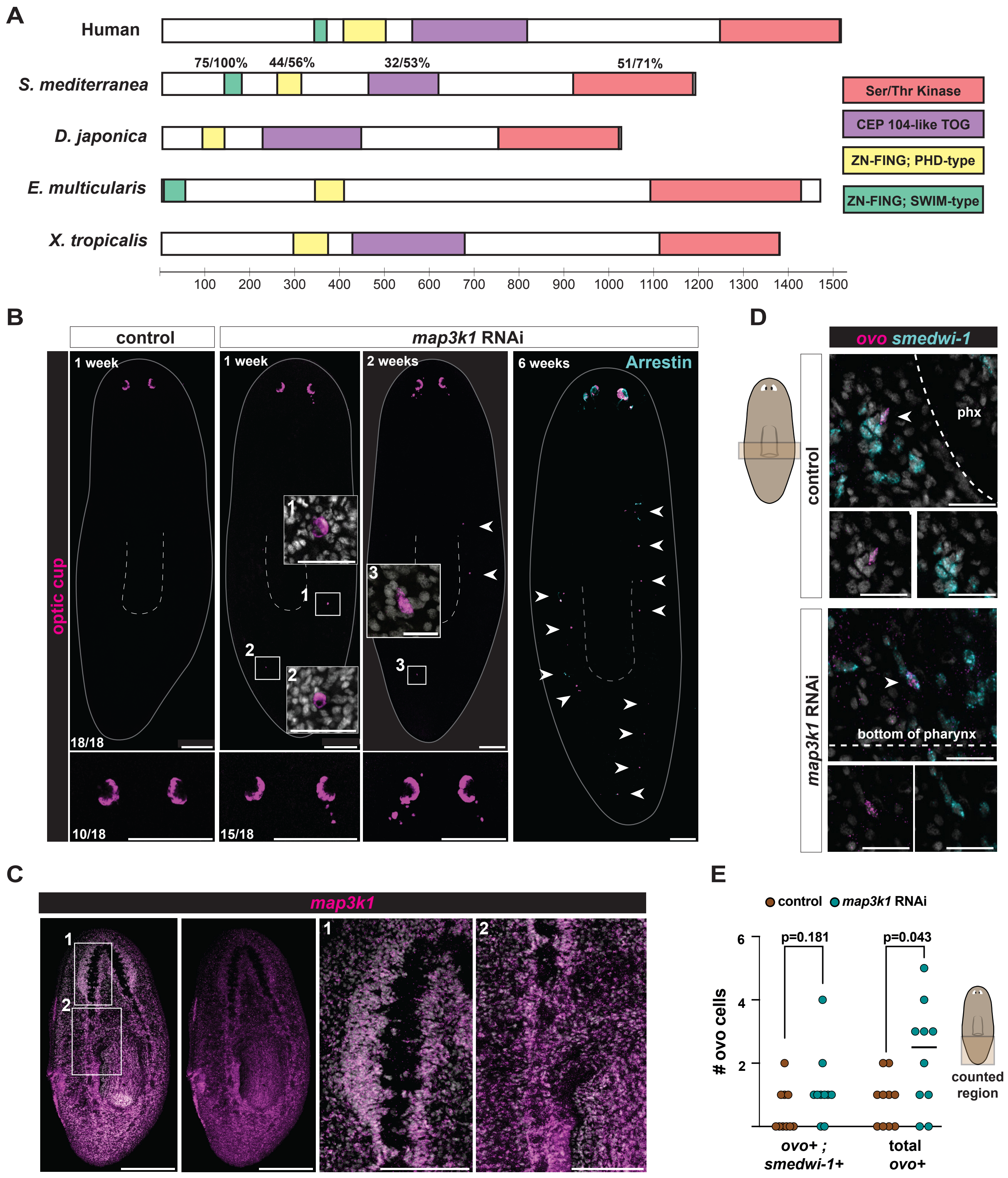

**Figure 1- figure supplement 1: *map3k1* RNAi results in differentiated eye cells throughout the AP axis, and no overt change in eye progenitor distribution**

**A.** Domain structure of MAP3K1 in human, *Schmidtea mediterranea, Dugesia japonica, Echinococcus multilocularis,* and *Xenopus tropicalis.* % identity/similarity is displayed above each domain for *Schmidtea mediterranea* Map3k1 compared to human MAP3K1. **B.** FISH images of *map3k1* RNAi animals at 1 and 2 weeks of *map3k1* RNAi showing ectopic OC cells (RNA probe pool to *catalase1, tyrosinase,* and *glut3*) in the head, trunk, and tail. The 1-week *map3k1* RNAi animal example included is from Figure 1C. Far right FISH shows ectopic OC and PRNs (anti-Arrestin) (white arrows) along the AP axis in a 6-week *map3k1* RNAi animal. Dorsal up. Scale bar, 200µm. **C.** FISH (using an RNA probe to *map3k1*) showing broad *map3k1* expression in a wild-type animal, with some visible expression in the brain and ventral nerve cords. Ventral, up. Scale bar, 200µm. **D.** FISH examples of *ovo*$^{+}$; *smedwi-1*$^{+}$ cells lateral to the posterior half of the pharynx in both *map3k1* RNAi and control animals after 3 weeks of RNAi. Dorsal up. Scale bar, 25µm. **E.** Quantification showing that *map3k1* RNAi animals have a similar albeit slightly higher total number of *ovo*$^{+}$ cells (*smedwi-1*$^{+}$ and *smedwi-1*$^{-}$) in the tail compared to control RNAi animals ($p=0.043$; Permutation test, 10,000 permutations; two-tailed); for comparison, *ovo*$^{+}$; *smedwi-1*$^{+}$ cells from Figure 1F are shown, which were not significantly different in the tail ($p=0.181$; Permutation test, 10,000 permutations; two-tailed – see Figure 1F). 3-week RNAi animals were used for counts.

### Figure 2- figure supplement 1

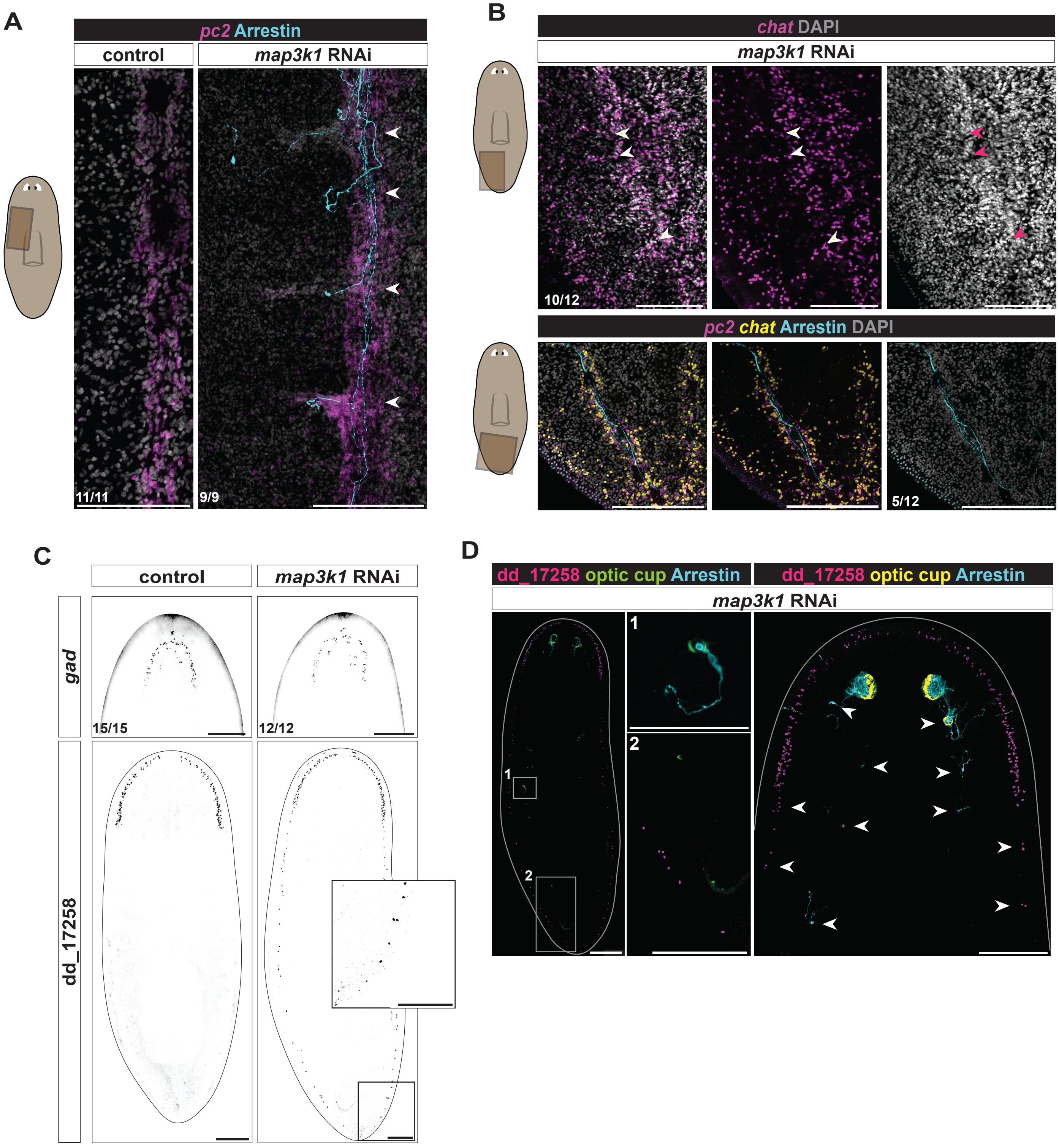

**Figure 2- figure supplement 1: *map3k1* RNAi results in posterior differentiation of some neural cell types**

**A.** Left panels: FISH images of the trunk regions of RNAi animals showing, in the case of *map3k1* RNAi, PRN (anti-Arrestin) projections along ventral nerve cord (VNC) and into ectopic brain branches (white arrows) labeled by DAPI and an RNA probe to the pan-neural marker, *pc2*. 5 weeks of RNAi. Scale bar, 100µm. Ventral up. Right panels: top row shows dorsolateral projections composed of *chat*$^{+}$ cells in the tail region of a *map3k1* RNAi (4 weeks RNAi) animal (n=10/12). Bottom row shows PRN projections (anti-Arrestin) within the VNC in the tail of a *map3k1* RNAi (5 weeks) animal (n=5/12). **B.** Top row FISH shows an example of brain branches (*chat*$^{+}$, white and pink arrows) in the tail of a *map3k1* RNAi animal (4 weeks RNAi). Bottom row FISH shows an example of PRN projections running along the VNC in the tail of a *map3k1* RNAi animal (5 weeks RNAi). Ventral up. Scale bars, 100µm. **C.** FISH showing *map3k1* RNAi animals (3 weeks RNAi) do not exhibit expansion of ventral *gad*$^{+}$ brain neurons, a stark contrast to ectopic dd_17258$^{+}$ cell differentiation along the entire AP axis (n=12/12). Images of dd_17258$^{+}$ neurons are full body perspectives of Figure 2C animals. Dorsal up. Scale bars, 200µm. **D**. FISH example of an animal with ectopic (arrows) OC cells (RNA probe pool to *catalase1, tyrosinase,* and *glut3*), photoreceptors (labeled with an anti-Arrestin antibody), and dd_17258$^{+}$ neurons extending down the AP axis. 5 weeks of RNAi. Dorsal up. Full body image scale bar, 200µm; all other panels scale bars, 100µm.

### Figure 3- figure supplement 1

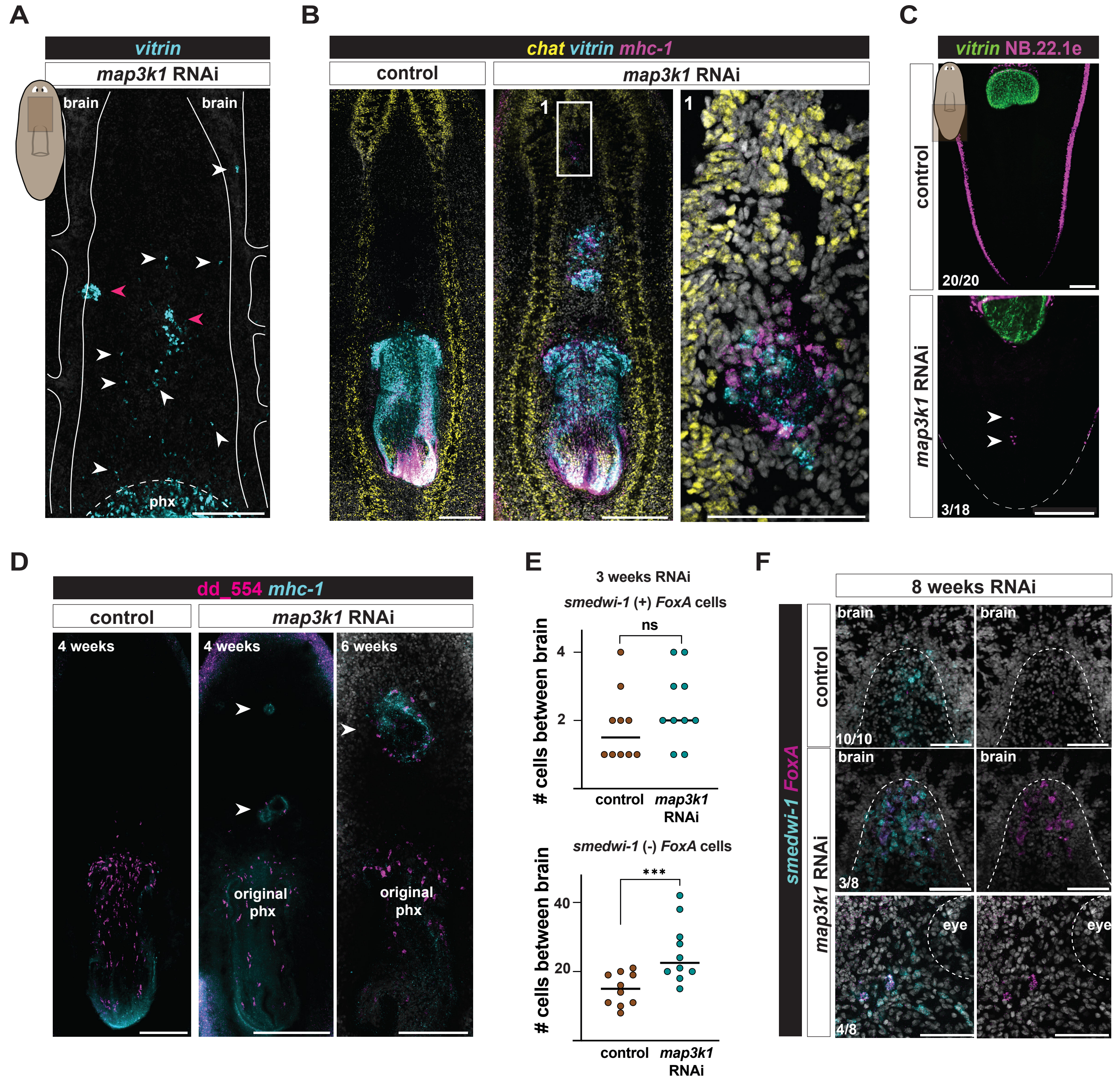

**Figure 3- figure supplement 1: *map3k1* RNAi causes ectopic differentiation of pharyngeal cell types**

**A.** Example FISH image of *vitrin*$^+$ cells (white arrows) and clusters of cells (pink arrows) anterior to the pharynx and in the ventral nerve cords in a 3-week *map3k1* RNAi animal. Ventral up. Scale bar, 100µm**. B.** FISH of a *map3k1* RNAi animal with clusters of *vitrin*$^+$ and *mhc-1*$^+$ cells in the pre-pharyngeal area, between the cephalic ganglia (*chat*$^+$). Animals from 6 and 8 weeks of RNAi were used. Ventral up. Scale bar, 200µm; magnified image scale bar, 50µm. **C.** FISH showing (n=3/18) *map3k1* RNAi animals have an ectopic focus (white arrows) of NB.22.1e$^+$ cells in the tail after 3 weeks of RNAi. Ventral up. Scale bar, 200µm. **D.** FISH image showing anterior clusters (white arrows) of dd_554$^+$ and *mhc-1*$^+$ cells in *map3k1* RNAi animals after 4 weeks and 6 weeks of RNAi. Ventral up. Scale bars, 200µm**. E.** Top graph shows no significant differences ($p=0.356$; Poisson regression) in the number of *FoxA*$^+$; *smedwi-1*$^+$ cells between the cephalic ganglia in *map3k1* RNAi and control animals after 3 weeks of RNAi. Bottom graph shows significantly more (***$p<0.0002$; Negative binomial regression) *FoxA*$^+$; *smedwi-1*$^-$ cells between the cephalic ganglia in *map3k1* RNAi compared to control animals. **F.** Some *FoxA*$^+$; *smedwi-1*$^+$ cells can be observed in the head of *map3k1* RNAi animals after 8 weeks of RNAi. Ventral up. Scale bar, 50µm. Numbers in each panel indicate number of animals displaying the result shown in the image out of total animals observed.

### Figure 4- figure supplement 1

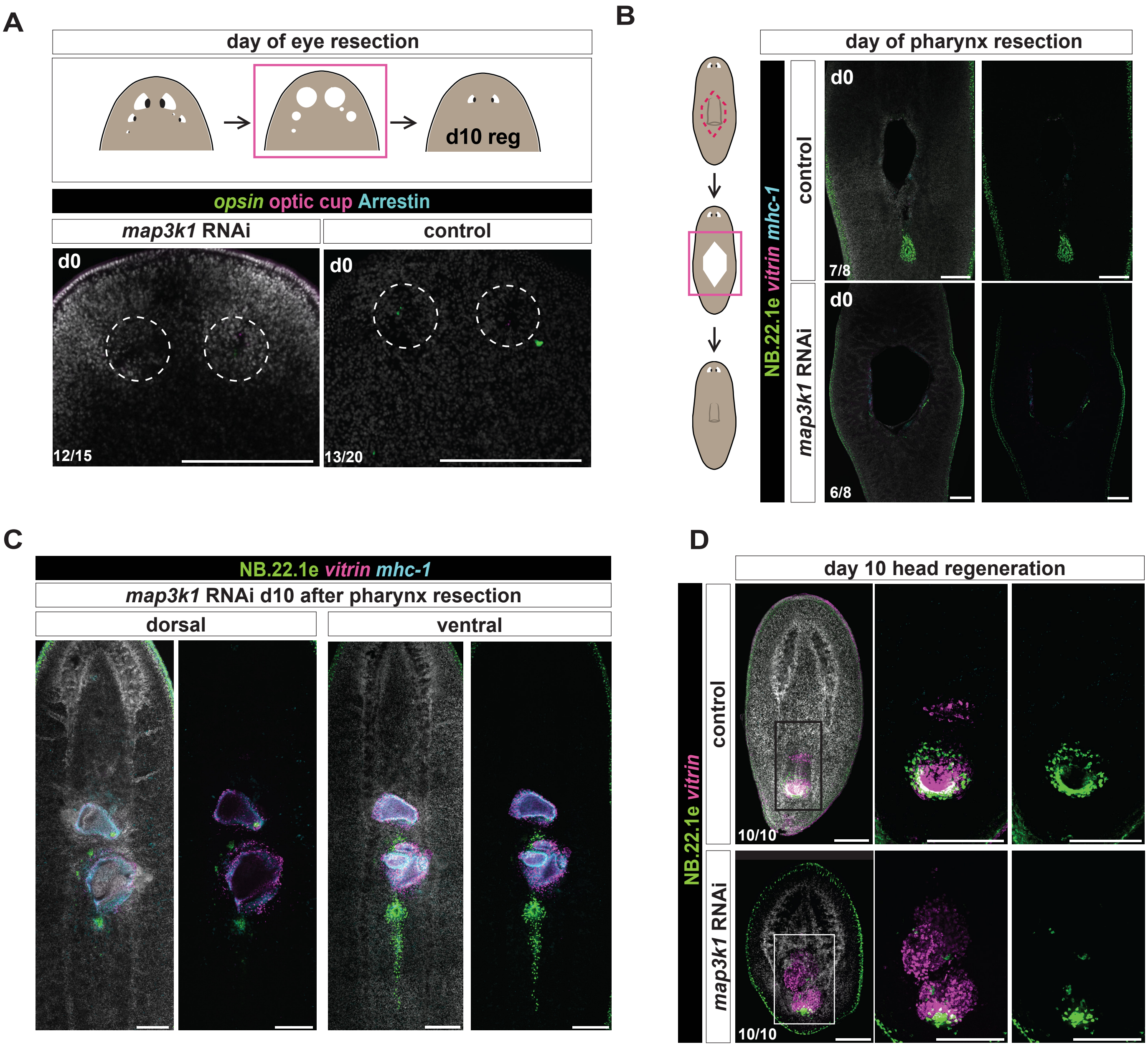

**Figure 4- figure supplement 1: *map3k1* RNAi animals undergo tissue-specific and whole-body regeneration, with some errors in pharynx regeneration**

**A.** FISH following eye resection at day zero (d0) with the markers for OC (*catalase1, tyrosinase, glut3* probe pool) and PRNs (*opsin* RNA probe, anti-Arrestin). 3 weeks of RNAi prior to resection. Scale bars, 200μm. Dorsal up. **B.** FISH following pharynx resection at day zero (d0) with the markers for *vitrin* (pharynx-specific), NB.22.1e (mouth and esophagus), and *mhc-1* (pharynx muscle). 3 weeks of RNAi prior to resection. Scale bars, 200μm. **C.** FISH example of *map3k1* RNAi animal 10 days following pharynx resection. Dorsal (left) and ventral (right) views of a disorganized pharynx regenerating in a *map3k1* RNAi animal. **D.** FISH of a *map3k1* RNAi head 10 days post-amputation showing a disorganized pharyngeal structure labeled with *vitrin* and NB.22.1e RNA probes. 3 weeks of RNAi prior to amputation. Scale bars, 200μm. Ventral up. Numbers in each panel indicate number of animals displaying the result shown in the image out of total animals observed.

### Figure 5 - figure supplement 1

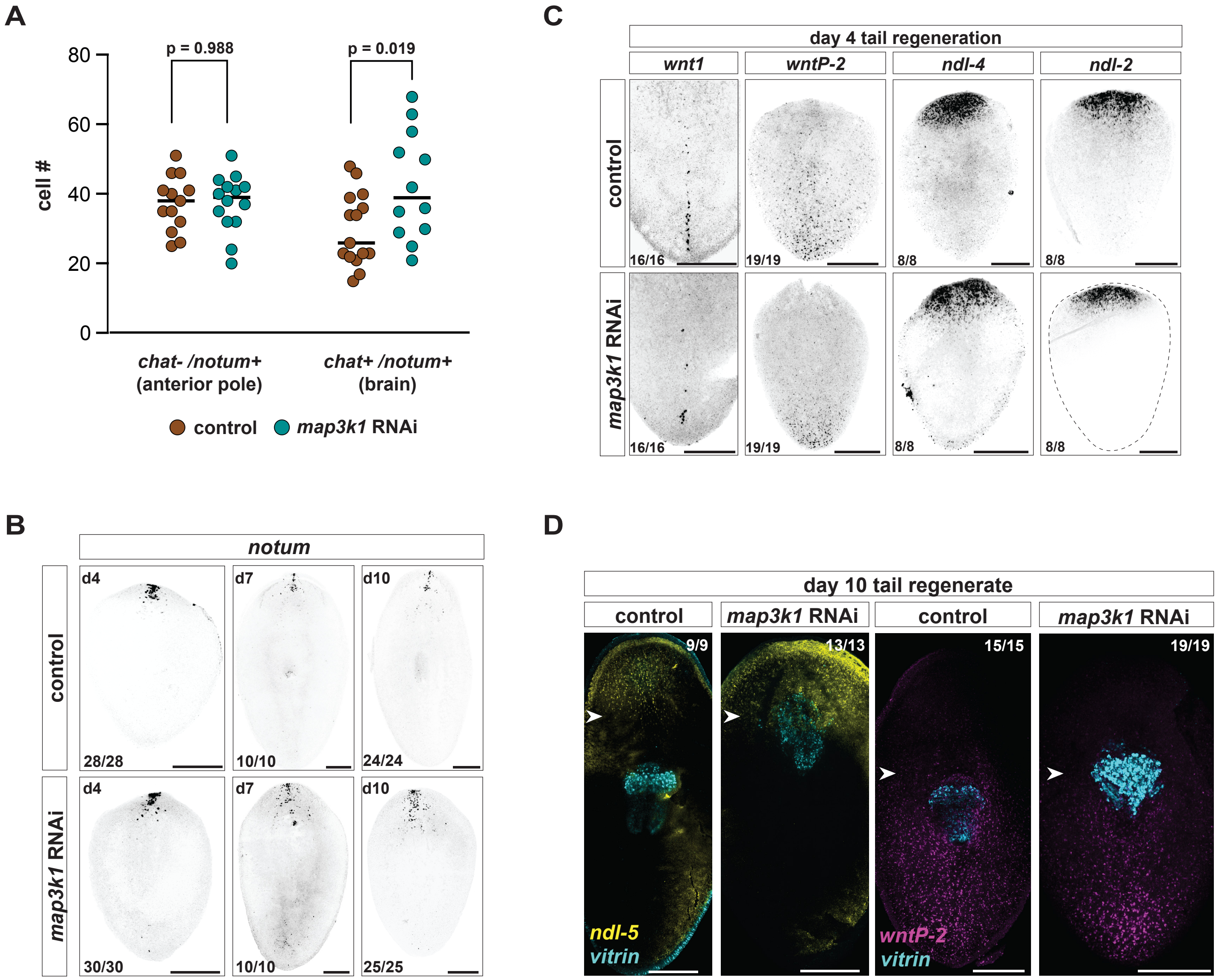

**Figure 5- figure supplement 1: *map3k1* RNAi tail fragments can regenerate the anterior pole but regenerate their new pharynx at a more anterior location**

**A.** Quantification showing no significant difference in pole *notum* cells (*notum*$^{+}$; *chat*$^{-}$) (p=0.988; Student's *t*-test) or brain *notum* cells (*notum*$^{+}$; *chat*$^{+}$) (p=0.027; unpaired Student's *t*-test) in *map3k1* RNAi animals compared to control animals. Animals fixed between 3-4 weeks of RNAi. **B.** FISH showing *notum* distribution in tail fragment at d4, d7, and d10 of regeneration. **C.** FISH showing *wnt1, wntP-2*, *ndl-4*, and *ndl-2* expression in d4 tail regenerates. Ventral up. Scale bars, 100µm. **D.** FISH showing pharynges regenerating inside of the *ndl-5* expression region*,* and outside of the *wntP-2* expression region in d10 *map3k1* RNAi tail regenerates. White arrows point to top of pharynx. Scale bar, 100µm. Ventral up. **B, C, D.** 3 weeks of RNAi occurred prior to amputations. Sample numbers are indicated in each panel.

### Figure 5- figure supplement 2

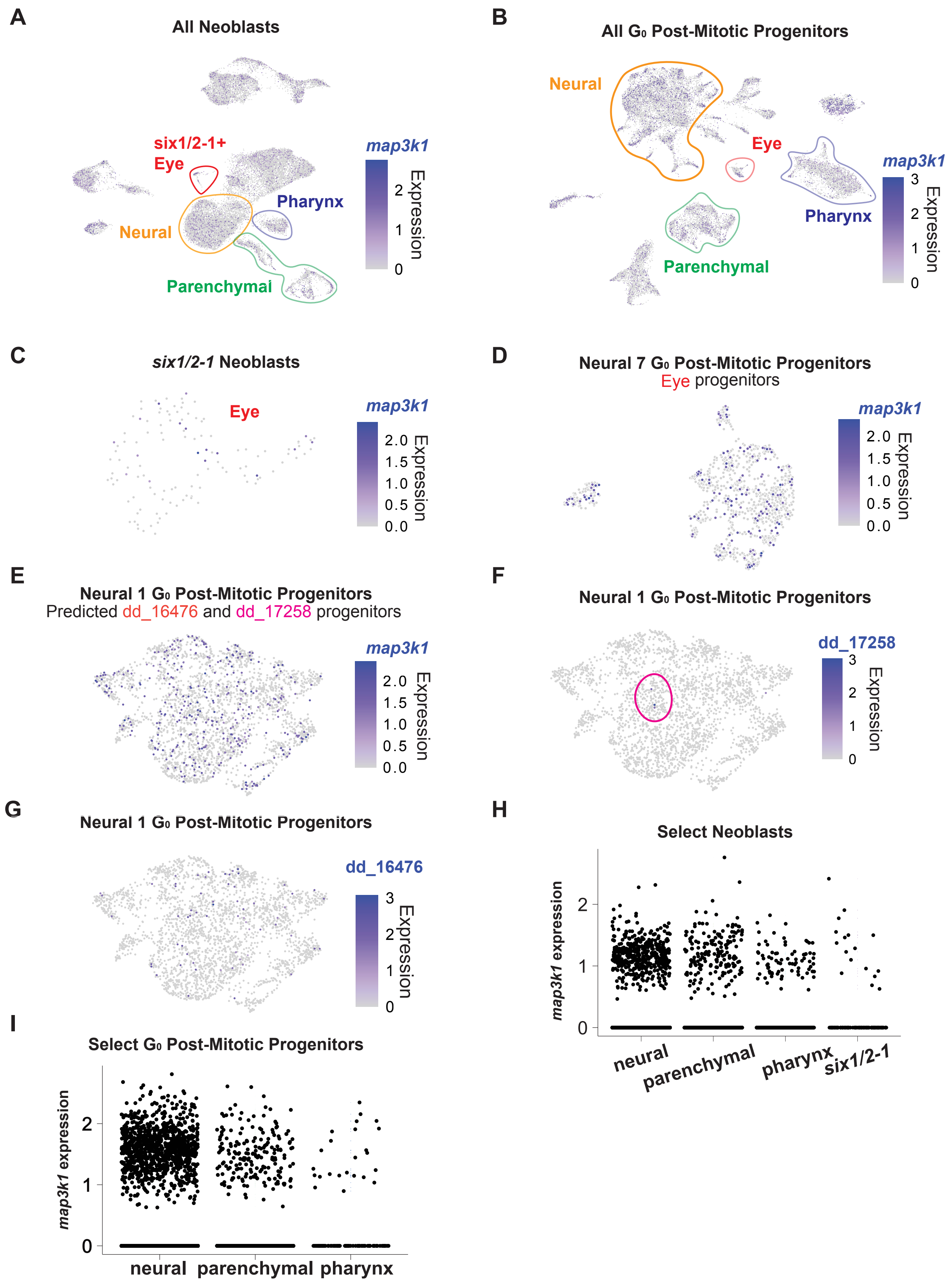

**Figure 5- figure supplement 2: *map3k1* is expressed in neoblasts and post-mitotic progenitors**

**A.** *map3k1* is expressed broadly across neoblast clusters. **B.** *map3k1* is expressed broadly across post-mitotic progenitors. **C.** *map3k1* is expressed in some *six-1/2-1*$^{+}$ neoblasts, which contain eye neoblasts. **D.** *map3k1* is expressed in some G0 post-mitotic eye progenitors (Neural 7 G0). **E.** *map3k1* is expressed within the predicted progenitor cluster (Neural 1) of the affected cell types: dd_17258 and dd_16476. **F.** dd_17258 mature marker expression within the predicted corresponding progenitor cluster (Neural 1 G0). **G.** dd_16476 mature marker expression within the predicted corresponding progenitor cluster (Neural 1 G0). **H.** *map3k1* expression in neural, parenchymal, pharyngeal, and *six1/2-1*$^{+}$ neoblast clusters. **I.** *map3k1* expression in neural, parenchymal, and pharyngeal post-mitotic progenitor clusters. **A, B, C, D, E, F, G, H, I.** All plot points are ordered.

### Figure 6 - figure supplement 1

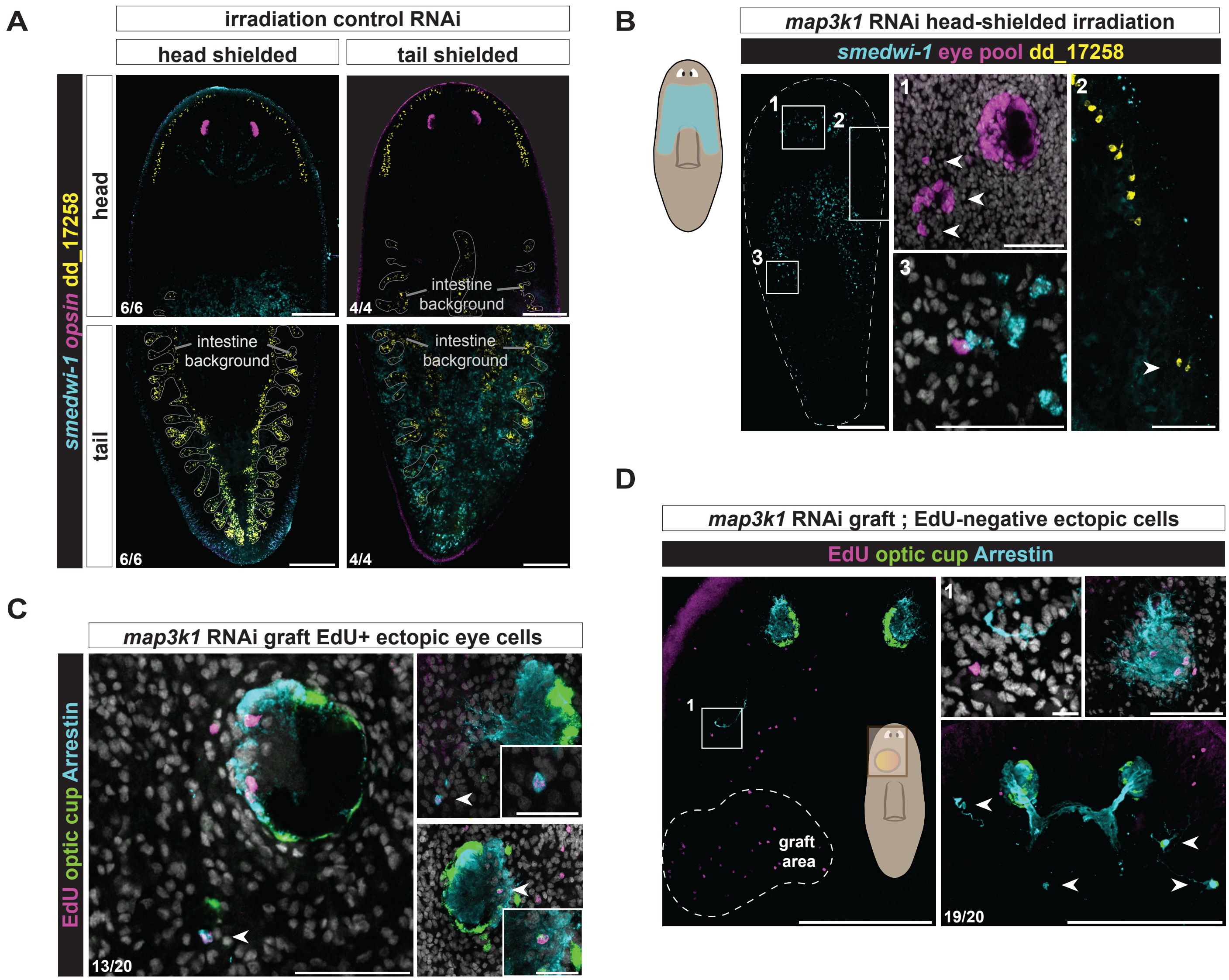

**Figure 6- figure supplement 1: *map3k1* RNAi animals display ectopic differentiation of local neoblasts**

**A.** FISH examples of head-shielded and tail-shielded irradiated animals that were fed control dsRNA showed no ectopic photoreceptors (*opsin*) or dd_17258$^+$ neurons in the head or tail region of either condition. See Figure 6A, B for experiment details. Scale bars, 200µm. Dorsal up. Yellow intestinal background signal is noted. *smedwi-1* labeling in a zone between the eyes and the pharynx is weak; this region frequently shows poor probe labeling across experiments, presumably associated with mucus production. **B.** An additional FISH example of a head-shielded, irradiated animal with ectopic eye cells (pool of RNA probes for *opsin*, *catalase1, tyrosinase, and glut3*) and dd_17258$^+$ neurons (white arrows for both ectopic cell types) in the region corresponding to live neoblasts (*smedwi-1*). Scale bars, 200µm; magnified images, 1-3 scale bars, 50µm. Dorsal up. **C.** FISH examples of EdU-positive ectopic eye cells (white arrows) in wild-type recipient animals that received a plug from an EdU-soaked *map3k1* RNAi animal. Scale bars, 20µm. Dorsal up. Two examples of EdU-labeled ectopic photoreceptors (anti-Arrestin) outside of the eye and one example of an OC cell (*catalase1/ tyrosinase*/*glut3* RNA probe pool) in the wrong place within the eye of a *map3k1* RNAi-plug recipient animal. **D.** FISH image of EdU-negative ectopic eye cells (white arrows) in wild-type recipient animals that received a plug from an EdU-soaked *map3k1* RNAi animal. Scale bar, 200µm; panel 1 scale bar, 10µm; upper right panel scale bar, 50µm. Dorsal up. **C, D.** Donors were fed between 2 and 3 weeks of dsRNA prior to transplantation. Numbers in each panel indicate number of animals displaying the result shown in the image out of total animals observed.

### Figure 7 - figure supplement 1

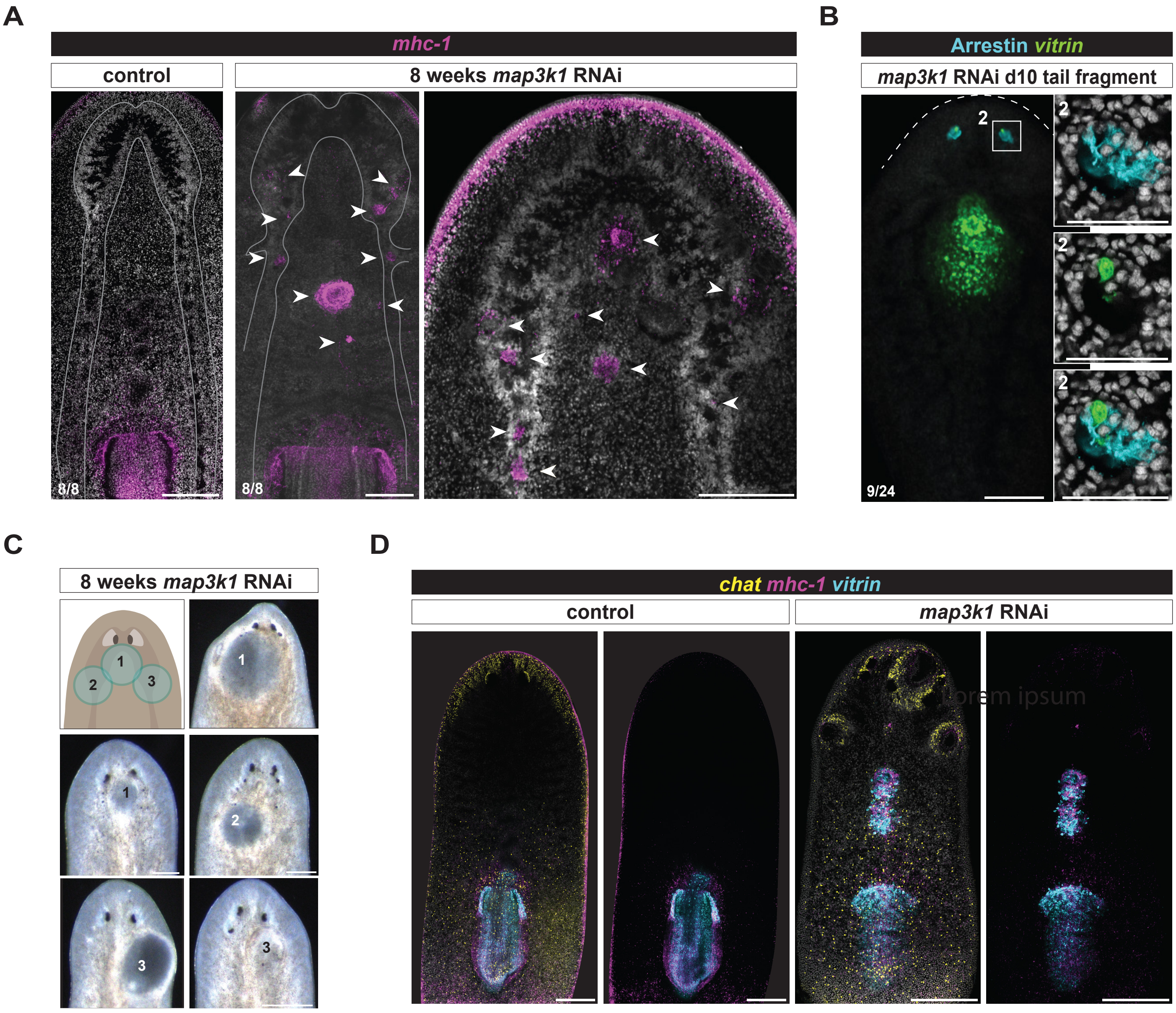

**Figure 7- figure supplement 1: *map3k1* RNAi animals display ectopic teratoma-like growths**
**A.** FISH showing a zoomed-out view of the ectopic *mhc-1* cells (white arrows) in the brain and ventral nerve cords of a *map3k1* RNAi animal (8 weeks RNAi) from Figure 7A. Scale bars, 200μm. Ventral up. **B.** Day 10 *map3k1* RNAi tail regenerate from Figure 7B with separated channels to show the ectopic *vitrin*$^{+}$ cell in the eye is not positive for the photoreceptor marker, Arrestin. Three weeks of RNAi occurred prior to fixation. Scale bar, 200μm. Zoom in scale bars, 50μm. Dorsal, up. **C.** Live image examples of teratomas after 8 weeks of *map3k1* RNAi. Scale bar, 200μm. Dorsal, up. **D.** FISH images showing a broader region of the animal in Figure 7D, and control RNAi; the *map3k1* RNAi animal shows *chat*$^{+}$*; mhc-1*$^{+}$ outgrowths. Animals were fed dsRNA for 6 to 8 weeks prior to fixation. Scale bar, 200μm; Dorsal, up.
